## Supplementary Information for "Synthetic Eco-Evolutionary Dynamics in Simple Molecular Environment"

### Synthetic Eco-Evolutionary Dynamics in Simple Molecular Environment. Supplementary Information

[a]Dipartimento di Biotecnologie Mediche e Medicina Traslazionale, Università degli Studi di Milano, Via Fratelli Cervi, 93 - L.I.T.A., Segrate, 20054, Italy; [b]Dipartimento di Fisica e Astronomia, Università degli Studi di Padova, Via Marzolo 8, Padova, 35131, Italy; [c]Department of Biomedical Sciences, Humanitas University, Via Rita Levi Montalcini 4, Pieve Emanuele, 20072, Italy; [d]IRCCS, Humanitas Clinical and Research Center, Via Manzoni 56, Rozzano, 20089, Italy

#### Tables with oligonucleotides sequence details

| | |
|---|---|
| Oligo1 | 5′-GGATGGGAGTGCTCTTCTTGAACTC-50N-AACTGCCTGGTGATACGACGATCGT-3′ |
| Blocker1-5′ | 5′-GAGTTCAAGAAGAGCACTCCCATCC-3′ |
| Blocker1-3′ | 5′-ACGATCGTCGTATCACCAGGCAGTT-3′ |
| Primer1-F | 5′-GGATGGGAGTGCTCTTCTTG-3′ |
| Primer1-R | 5′-ACGATCGTCGTATCACCAG-3′ |
| Resource1 | 5′- /5AmMC12/CGGTATTGGACCCTCGCATG-3′ |
| Oligo2 | 5′-CCCTATGCGACCCTCCGATGTAGAC-50N-CCTTGAGATTGCCGATCCATCCTCG-3′ |
| Blocker2-5′ | 5′-GTCTACATCGGAGGGTCGCATAGGG-3′ |
| Blocker2-3′ | 5′-CGAGGATGGATCGGCAATCTCAAGG-3′ |
| Primer2-F | 5′-CCCTATGCGACCCTCCG-3′ |
| Primer2-R | 5′-CGAGGATGGATCGGCAAT-3′ |
| Resource2 | 5′-/5AmMC12/CGTATCACCAGGCAGTTGAG-3′ |

**Table 1.** Sequences of the oligonucleotides used in this work.

| | |
|---|---|
| 13.0 | FS1-5′-ACCACGCCAAGACTTCTGACCATGCGAGGGTCCGTTACGGATGTCGATCGG-FS1-3′ |
| 11.0 | FS1-5′-CGTGCACTGAAAGGACGCGTCGTGGTAGGGGGACGTCATGCGAGGGTTCCT-FS1-3′ |
| 8.0 | FS1-5′-ACCGCGCATGGCCTTGACGTAGCGACGGTGTCTGTGGCGCGATGGAGGGCG-FS1-3′ |
| 14.0 | FS1-5′-ACTCGGCAGGTAGGCGGTCCCTTTGACATGCGAGGGTCCACGTCGGTGTGT-FS1-3′ |
| 7.0 | FS1-5′-TGCGGCGAGGGGTTCGGCGGCGAGTGGTCCGTGTCGTCGGATTGGCCTAAC-FS1-3′ |
| 12.0 | FS1-5′-ACTCGCCGCTGGTCGTGGTTGCGAGGGTCCATCCCTAGTTCAGAGCGTTGG-FS1-3′ |
| 12.1 | FS1-5′-ACTCGCCGTACGCGAACGCCCCGTCGTACGGTTTCATGCGAGGGTCGCGCG-FS1-3′ |
| 5.0 | FS1-5′-CGACAGCGGGGGGACAGTCACTGCGGGCATGCAGGGTGGTGCGGGCGTAAC-FS1-3′ |
| 12.2 | FS1-5′-ACTCGGCAGGTAGGCGGTCCCTTTGACATGCGAGGGTCGCGCGGGATGGGA-FS1-3′ |
| 5.1 | FS1-5′-TCGCGTCCAGTGTGGTGATGGGCAGTGTGGCTTTCGTGTGTAGGAACTGCC-FS1-3′ |

**Table 2.** Oligo1 cycle 24 top 10 sequences. 'FS1-5′' stands for 'fixed sequence 1 at 5′' and it similarly holds for 'FS1-3′'.

| | |
|---|---|
| 11.1 | FS1-5′-ACCTGCGAGGGTCCGTGGTTACGATTAGCGGGAGAACAGTGGCATGTCGGC-FS1-3′ |
| 7.X | FS1-5′-AATCGCAGCGAGGGGACAAGGCCGAAAATGGTTCCGGCGGAGTGGAACCAG-FS1-3′ |

**Table 3.** Oligo1 other relevant sequences. 'FS1-5′' stands for 'fixed sequence 1 at 5′' and it similarly holds for 'FS1-3′'.

#### Supplementary Figures of Fig. 1

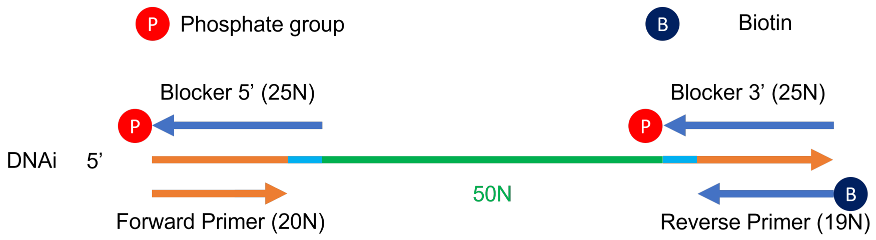

**Figure 1 - figure supplement 1.** Detailed structure of DNA individuals.

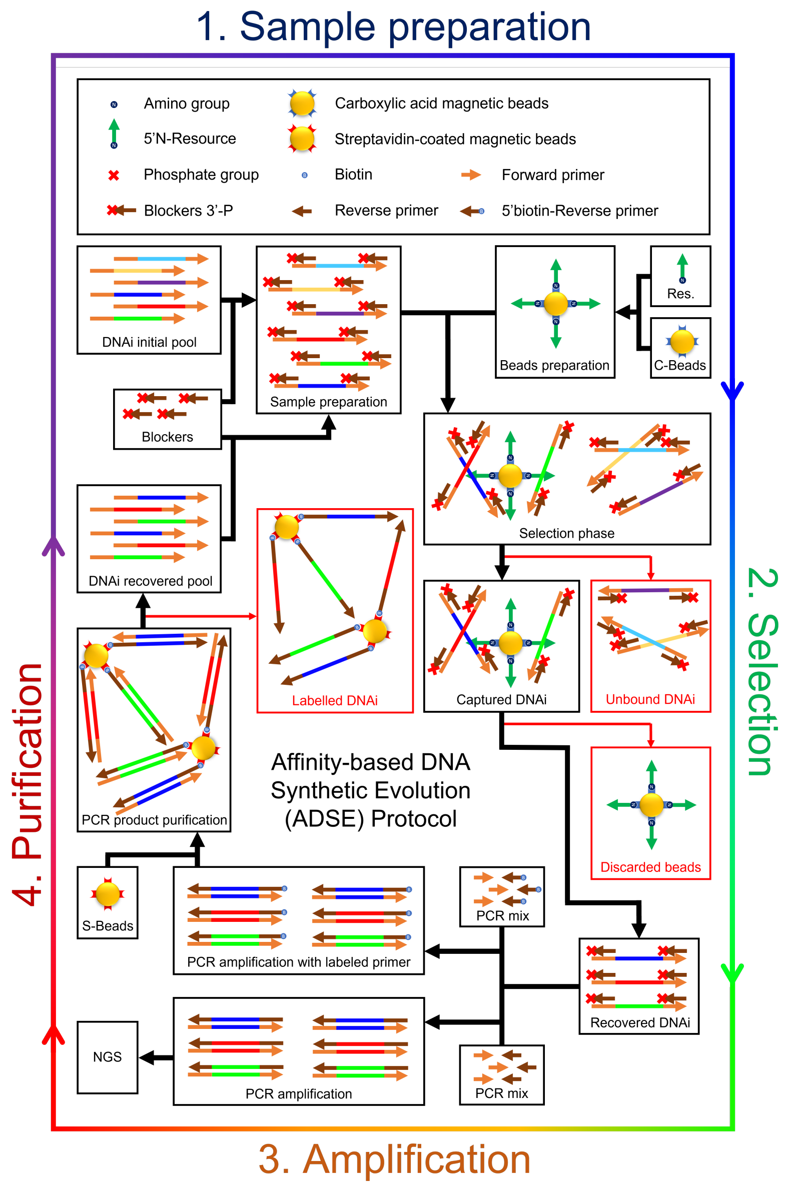

**Figure 1 - figure supplement 2.** Detailed scheme of the Affinity-based DNA Synthetic Evolution protocol.

### Supplementary Figures of Fig. 2

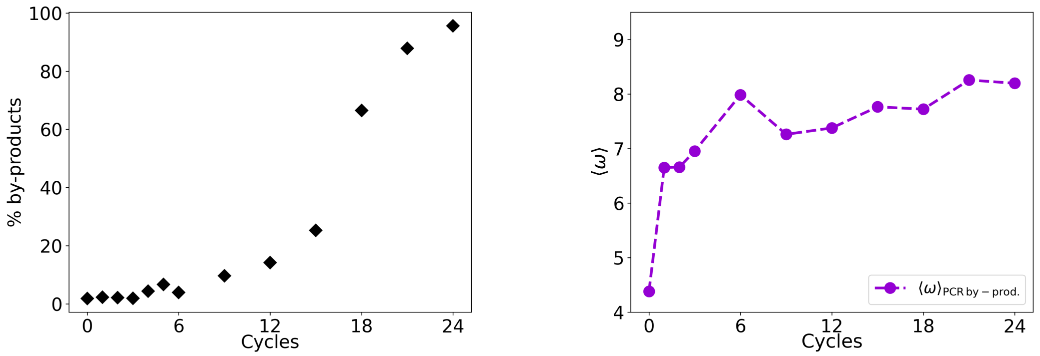

(a) (b)

**Figure 2 - figure supplement 1. a)** % of by-products as a function of the evolutionary cycles in the Oligo1 experiment; **b)** Evolution of $\langle \omega \rangle$ considering only PCR by-products.

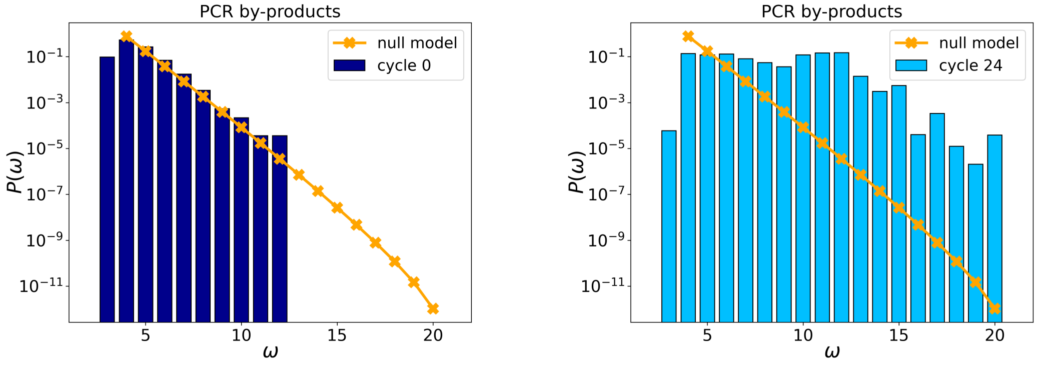

**Figure 2 - figure supplement 2.** probability distributions $P(\omega)$ of PCR by-products for at cycles 0 and 24. The orange points and line are the distributions evaluated with the null model.

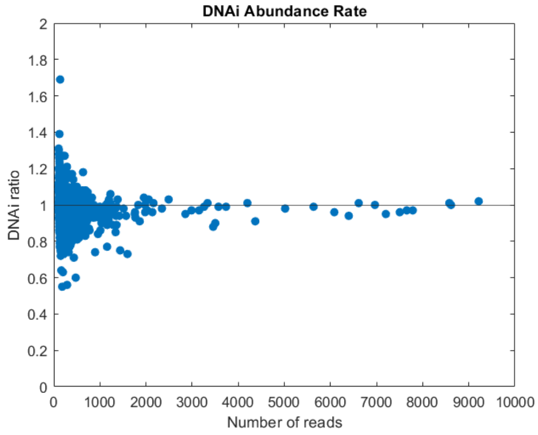

**Figure 2 - figure supplement 3.** Compared abundance of DNAi populations between two sequencing replicates of the same library (cycle 9 of oligo 1). The ratio is computed between the number of reads of each DNA species shared by the two sequencing runs after normalization for the total number of reads.

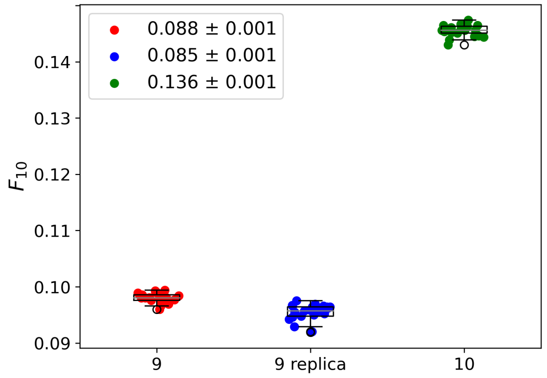

**Figure 2 - figure supplement 4.** Whisker plots showing the fraction of 10 most abundant individuals $F_{10}$ for cycle 9 (red), cycle 9 replica (blue) and cycle 10 (green). For each dataset, every data point represents one out of 20 samples of $4 \times 10^5$ sequences. To compare the three systems we extracted from each the fraction of the population formed by the 10 most abundant species, $\langle F_{10} \rangle$. The legend reports the average $\langle F_{10} \rangle \pm$ the standard deviation $\sigma$ (among the 20 subsamples) for each group of data points. This analysis leads to an average of 8.8% $\pm$ 0.1% of the population for cycle 9 (red dots), of 8.5% $\pm$ 0.1% for cycle 9 replica (blue dots) and of 13.6% $\pm$ 0.1% for cycle 10 (green dots). Cycle 9 and Cycle 9 replica are statistically compatible within 3 $\sigma$. The similarity between cycle 9 and cycle 9 replica and the marked difference between cycle 9 replica and cycle 10 indicates that the relevant part of the selection is indeed performed by the resource-binding mechanism, while drifts induced by PCR play a secondary role. As a further check, we compared the specific sequences across the 20 samples in cycle 9 and cycle 9 replica datasets and found that the 10 most abundant sequences are almost always the same. In particular, the first 8/9 are always the same, possibly shuffled.

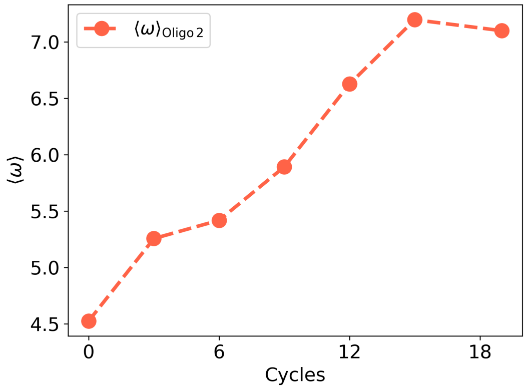

**Figure 2 - figure supplement 5.** $\langle \omega \rangle$ as a function of the experimental cycles for Oligo2.

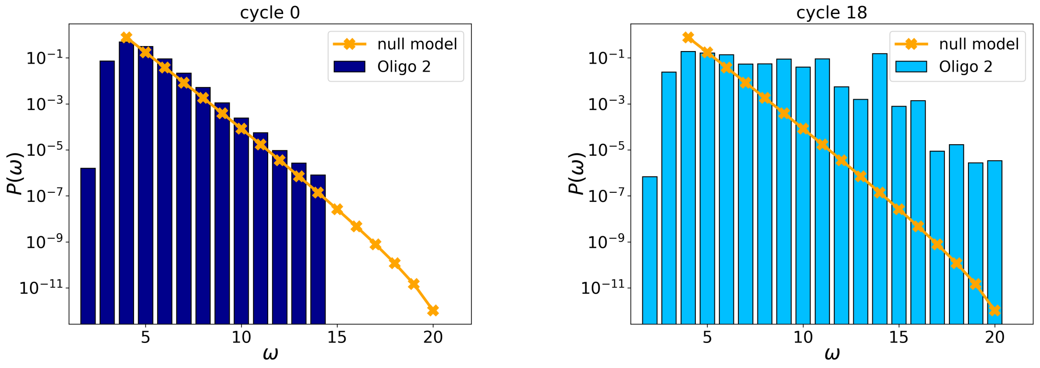

**Figure 2 - figure supplement 6.** Initial (left, cycle 0) and final (right, cycle 18) $p(\omega)$ distributions for Oligo2.

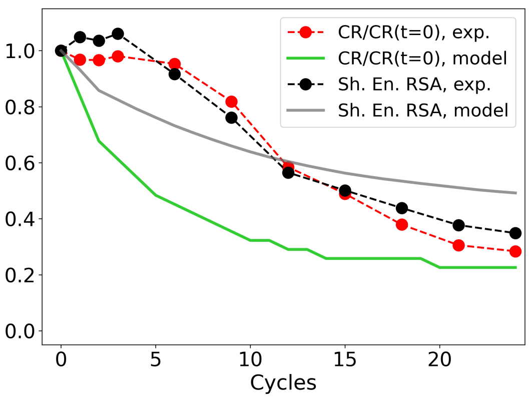

**Figure 2 - figure supplement 7.** evolution of the zip ratio CR for the file with the list of sequences (red dots), normalized by its value at time 0; the same for the IBEE model (green line); and of the Shannon Entropy associated to the RSA distribution (experimental: black dots, IBEE: gray line). The IBEE model has $\gamma = 3$, $\omega_{sat} = 10$, $10^6$ individual and $10^4$ resources and the data shown are the average of 20 statistically independent realizations.

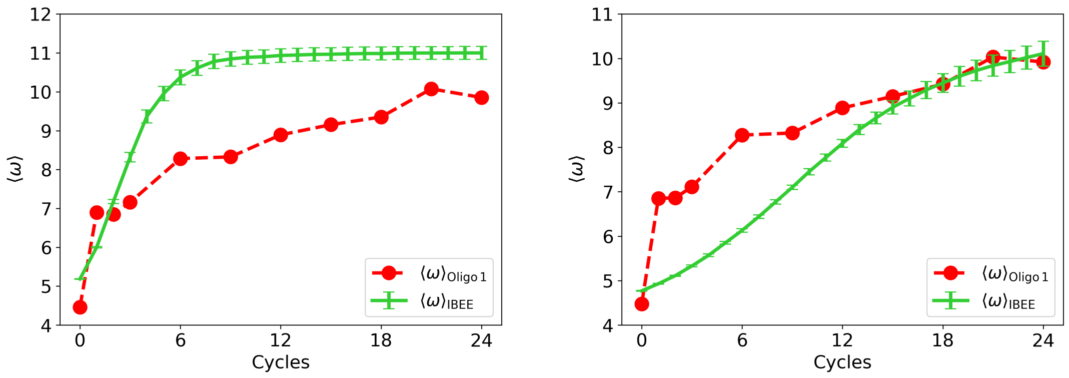

**(a)** **(b)**

**Figure 2 - figure supplement 8. a)**: $\langle\omega\rangle$ as a function of the experimental (red) and simulated (green) cycles. This IBEE model has a fitness $f(\omega) = \left(\frac{\omega}{\omega_{max}}\right)^{\gamma}$, with $\gamma = 3$ and without any saturation at $\omega_{sat}$. **b)**: $\langle\omega\rangle$ as a function of the experimental (red) and simulated (green) cycles. This IBEE model has a fitness $f(\omega) = \left(\frac{\omega^*}{\omega_{max}}\right)^{\gamma}$, with $\omega_{sat} = 10$ and $\gamma = 1$ for the whole simulation.

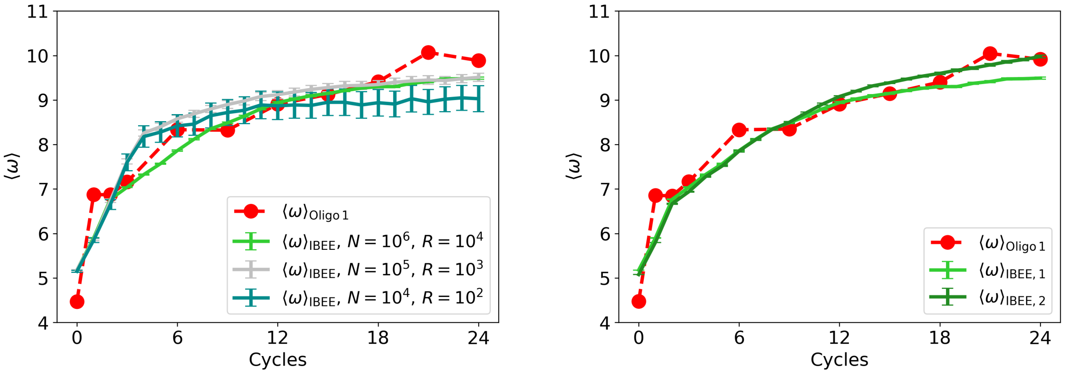

**(a)** **(b)**

**Figure 2 - figure supplement 9. a)**: $\langle\omega\rangle$ as a function of the experimental (red) and simulated cycles. Green: $N = 10^6,\ R = 10^4$; silver: $N = 10^5,\ R = 10^3$; teal: $N = 10^4,\ R = 10^2$. **b)**: $\langle\omega\rangle$ as a function of the experimental (red) and simulated cycles (IBEE model shown in the main text, #1: green. IBEE model #2 with a different starting population, dark green.)

#### Supplementary Figures of Fig. 3

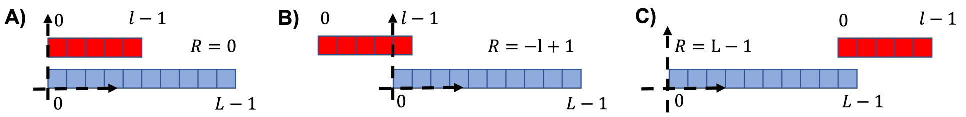

**Figure 3 - figure supplement 1.** Three examples of different relative positions for the attachment. **a)**: $R = 0$, i.e. the two sequences left ends are found in the same position. **b)**: leftmost possible position for the red sequence ($R = -l + 1$). **c)**: $R = L_1$, corresponding to the rightmost position of the red sequence.

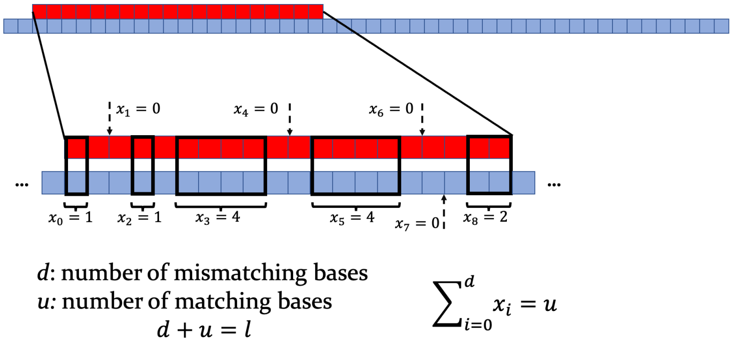

**Figure 3 - figure supplement 2.** Top: scheme of the target strand (red) and of a longer consumer strand (blue). Bottom: zoom on the interaction region, with $d$ mismatching nucleotides and $u$ matching ones. Note that one $x_i$ is computed between any pair of consecutive non overlapping bases. When $x_i = 0$, a dashed arrow indicates the middle point between the two pairs of mismatching bases involved.

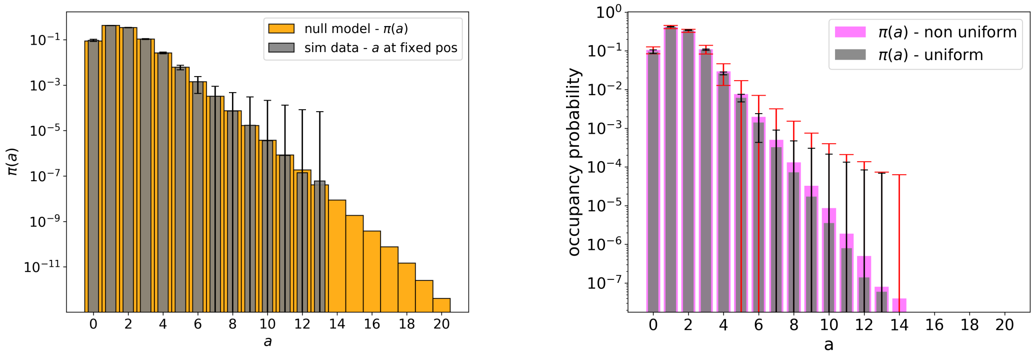

**(a)** **(b)**

**Figure 3 - figure supplement 3. a):** Orange: null model without threshold, analytic distribution $\pi(a)$. Grey: simulated null model, 50-mers population sample = $10^6$ individuals: mean over 50 repetitions and errorbars are reported. Note the log scale on vertical axis. **b)**: grey bars with black errorbars have the same meaning as in panel a). Pink: the same but with a non-even distribution of the 4 nucleotides (A=23%, C=19%, G=29%, T=29%) in the initial population, with red relative errorbars.

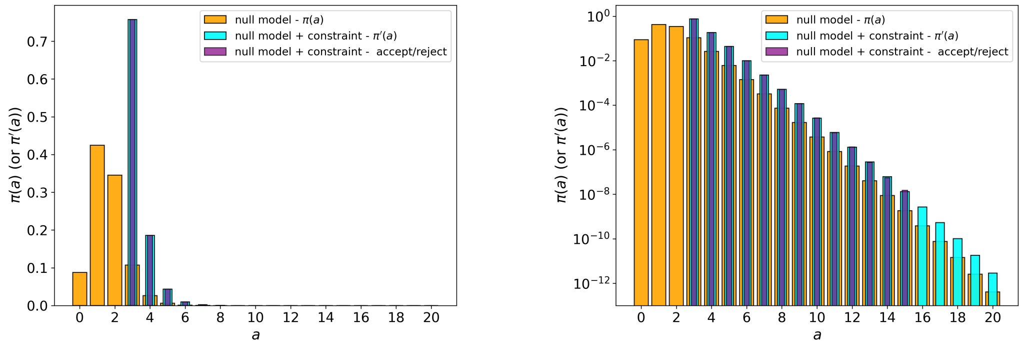

**(a)** **(b)**

**Figure 3 - figure supplement 4. a)**: Orange: null model without threshold, analytic $\pi(a)$. Cyan: null model with threshold on $a < T$ values, $T = 3$, analytic $\pi'(a)$. Purple: average over 200 runs of the null model with threshold on $a < T$ values, $T = 3$, simulated with explicit rejection of $10^6$ values. **b)**: same as panel a), but in log scale.

#### Supplementary Figures of Fig. 4

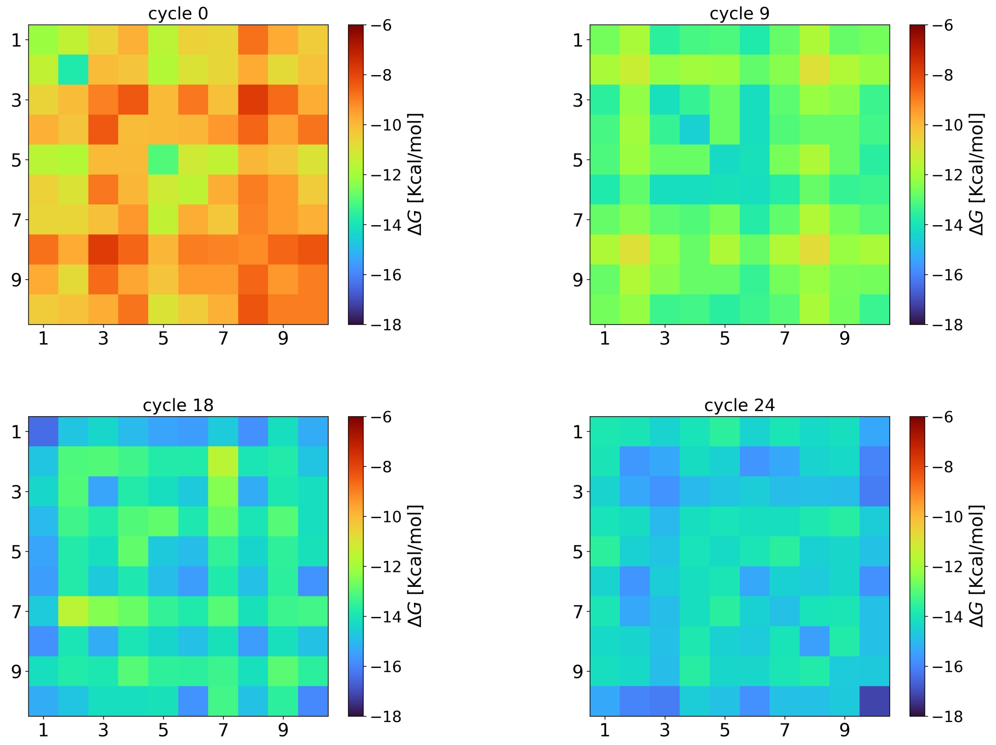

**Figure 4 - figure supplement 1.** Evolution of the intra-species interaction strengths ($\langle \Delta G_{pp} \rangle$). At cycle 0, the interactions strengths are compatible with those obtained from completely random strings ($-10.1 \pm 0.7$ kcal/mol). However, during the eco-evolutionary dynamics, the interactions strengths among the surviving strings increase (cycle 9: $-13.0 \pm 1.0$ kcal/mol; cycle 18: $-14.0 \pm 0.8$ kcal/mol) reaching the maximum values (on average) at the last cycle ($-14.8 \pm 0.9$ kcal/mol). This result indicates that there is a selection favoring species that can interact among them. Hyper-parameters used in these NUPACK calculations are the same described in the Methods section of the main text.

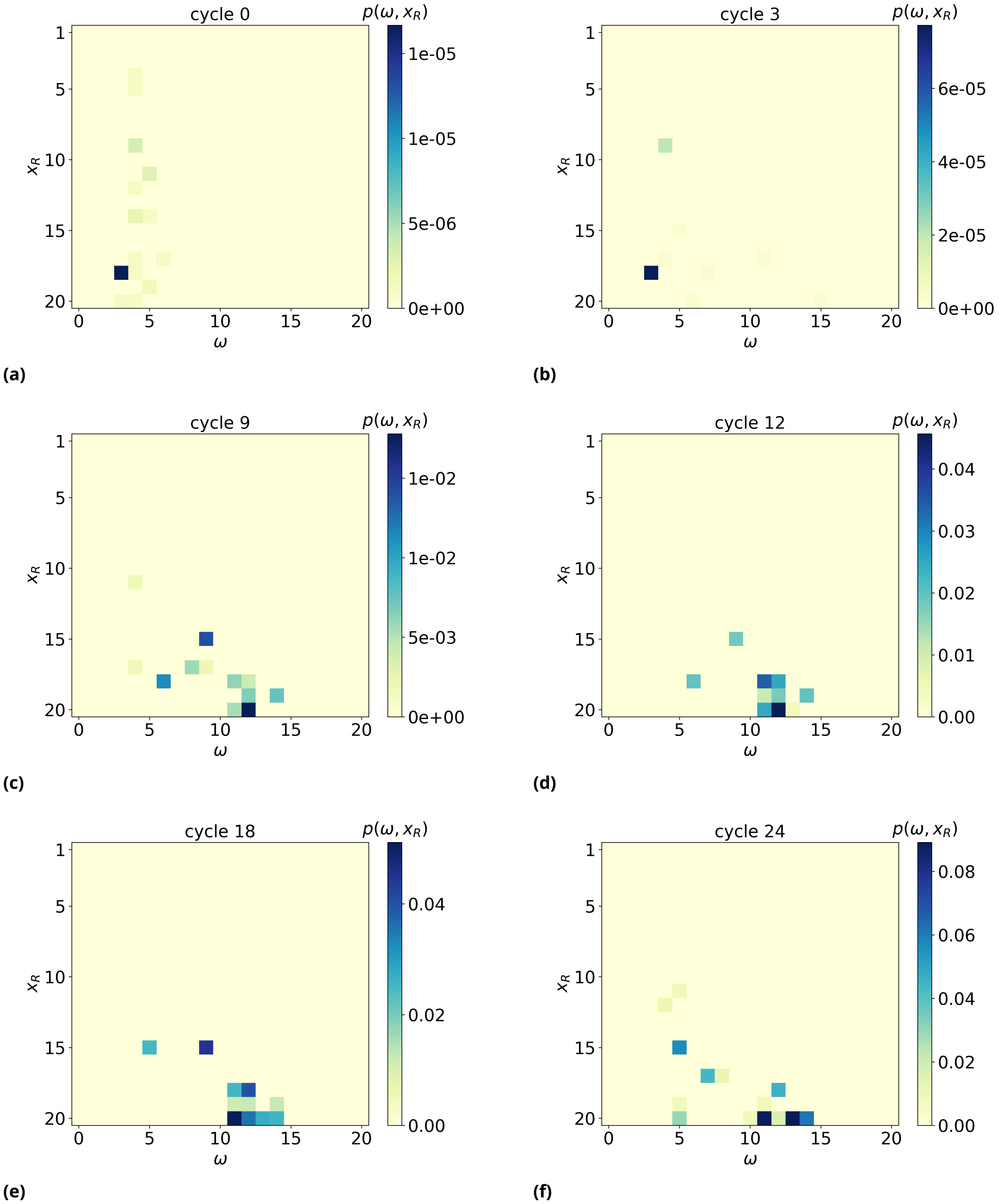

**Figure 4 - figure supplement 2.** 2D probability distribution $p(\omega, x_R)$, as a function of time (from top left to bottom right). We observe a significant shift of $x_R$, the rightmost basis of the of the resource involved in the formation of such an MCO, towards high values, meaning the MCO are preferably formed far away from the bead surface, near the free end of target strands.
